# Sperm competition favours intermediate sperm size in a hermaphrodite

**DOI:** 10.1101/2023.12.12.571300

**Authors:** Santhosh Santhosh, Dieter Ebert, Tim Janicke

## Abstract

Sperm competition is a potent mechanism of post-copulatory sexual selection that has been found to shape reproductive morphologies and behaviours in promiscuous animals. Especially sperm size has been argued to evolve in response to sperm competition through its effect on sperm longevity, sperm motility, the ability to displace competing sperm and ultimately fertilization success. Additionally, sperm size has been observed to co-evolve with female reproductive morphology. Theoretical work predicts that sperm competition may select for longer sperm but may also favour shorter sperm if sperm size trades off with number. In this study, we studied the relationship between sperm size and post-mating success in the free-living flatworm, *Macrostomum lignano*. Specifically, we used inbred isolines of *M. lignano* that varied in sperm size to investigate how sperm size translated into the ability of worms to transfer and deposit sperm in a mating partner. Our results revealed a hump-shaped relationship with individuals producing sperm of intermediate size having highest sperm competitiveness. This finding broadens our understanding of the evolution of sperm morphology by providing empirical support for stabilizing selection on sperm size under sperm competition.

## Introduction

Ever since Parker’s pioneering work on sperm competition, the study of post-copulatory sexual selection has drastically improved our understanding of male and female reproductive biology (Parker, 1970, 1982; Birkhead and Moller, 1999; Simmons, 2001). In species where individuals mate multiply with different partners, post-mating fertilisation success determined by the efficiency to outcompete rival ejaculates can become a major predictor of male reproductive success (Collet et al., 2012; Pélissié et al., 2014; Marie-Orleach et al., 2021). Consequently, sperm competition is expected to promote post-copulatory intrasexual selection on various traits that influence fertilization success. Over the last decades, great efforts have been made to gain a better understanding of such phenotypic effects of ejaculate traits on the outcome of sperm competition. Especially, sperm size has received substantial attention (Simmons and Siva-Jothy, 1998; Simmons,2001; Pizzari and Parker, 2009) and has been documented to have highly variable relationships (positive, negative and no association) with sperm motility, sperm longevity and the defensive behaviours of an ejaculate across species (Snook 2005; Fitzpatrick and Lüpold 2014). In addition, sperm size has been found to vary tremendously not only within ejaculates but also among males and among species, which is often considered to result from selection arising from sperm competition and/or cryptic female choice (Pitnick et al., 2009). Theoretical work often assumes a positive effect of sperm size on its fertilization success under sperm competition (Parker, 1993; Parker et al., 2010; Pizzari and Parker, 2009). Yet, sperm size is also considered to trade-off with sperm number, which complicates the prediction of how sperm size is selected under varying sperm competition risks and intensities. Specifically, if sperm size trades-off against the number of sperm produced or resources required for pre-copulatory mate acquisition, selection arising from sperm competition may favour ejaculates with sperm of intermediate size (Parker, 1993; Parker et al., 2010; Parker and Begon, 1993).

Although variation in ejaculate performance traits, such as sperm size, is generally intrinsic to males, it has also been shown that females play an active and crucial role in post-copulatory sexual selection and that competitive fertilization success of males is influenced by interactions between sperm and the female reproductive tract (Lüpold et al., 2013; Miller and Pitnick, 2002; Pitnick et al., 2009). Notably, sperm size has been found to covary with the length of female reproductive organs, such as seminal receptacle or spermatheca (Minder et al., 2005; Morrow and Gage, 2000; Pitnick et al., 1999, 2003; Briskie and Montgomerie 1992, Briskie at al., 1997). Moreover, using an experimental evolution study, Miller and Pitnick (2002) showed a correlated evolution in sperm length as a result of an evolutionary increase in seminal receptacle length in *Drosophila melanogaster*. Thus, selection on sperm size may also arise from selection imposed by properties of the female reproductive tract.

Empirical tests on the relationship between sperm size and sperm competitiveness provided mixed results within and across species (Pizzari and Parker, 2009). Comparative studies indicate that higher levels of sperm competition are associated with longer sperm in internally fertilizing species (e.g., fish: Balshine et al., 2001, birds: Briskie et al., 1997, Lepidoptera: Gage 1994, mammals: Gomendio and Roldan, 1991 & Tourmente et al., 2011; but also see, Stockley et al., 1997) suggesting that sperm size is determined, at least partly, by the intensity of sperm competition. By contrast, negative or no relationship between sperm competition and sperm size has been reported for Sylvian warblers (Immler and Birkhead, 2007) and mammals (Gage and Freckleton, 2003). Another line of across-species comparisons suggests that higher levels of sperm competition are associated with reduced intra-specific and intra-male variation in sperm size across birds (Calhim et al., 2007, Immler et al., 2008; Kleven et al., 2008), and such results have been speculated to arise from stabilizing selection for an optimum sperm size (Calhim et al., 2007; Lüpold et al., 2009).

The most direct evidence for selection acting on sperm size comes from studies quantifying how phenotypic variation in sperm size translates into sperm competitive success within species. Empirical studies attempting to establish such associations within species have found positive (Radwan, 1996), negative (Gage and Morrow, 2003; Garcìa-González and Simmons, 2007) or no clear relationship (Cascio Sætre et al., 2018; Kahrl et al., 2021) between sperm size and male reproductive success (or paternity share). In addition, studies comparing relative fertilization success between extreme sperm sizes (e.g. large vs small) also provided mixed results (Bennison et al., 2015; Morrow and Gage, 2001). These contrasting results might, at least partly, result from substantial within-individual variation in male post-mating success and the associated uncertainty in estimating individual differences in sperm competitiveness. Marie-Orleach et al. (2021) took measures of reproductive performance of the same individual over multiple mating events and detected low repeatabilities for different proxies of post-mating success, indicating the significance of stochastic variation in measures of post-mating success. Importantly, the vast majority of empirical studies assume a linear relationship between sperm size and reproductive (or post-mating) success and do not test for non-linear associations which limits our understanding on how selection operates on sperm size. To our knowledge, only five studies that have tested for such non-linear associations, but none of the studies found any significant non-linear associations between sperm length and estimates of fitness (Cramer et al., 2013; Lymbery et al., 2018; Kahrl et al., 2021; Rowe et al., 2022; Marie-Orleach et al., 2023).

Here, we report an experimental study exploring the relationship between sperm size and post-mating success within a species. Specifically, we tested the hypothesis that sperm size predicts sperm transfer success in the simultaneously hermaphroditic flatworm, *Macrostomum lignano,* a species in which sperm competition has been shown to be intense (Janicke et al., 2013; Marie-Orleach et al., 2021). In *M. lignano*, a recent study explored how sperm morphology predicts male reproductive success (and other fitness components) (Marie-Orleach et al., 2023), focussing on purely phenotypic relationships between sperm traits and male fitness components. By using inbred lines, we aimed to extend previous findings by exploring how sperm competition generates selection on standing genetic variation in sperm length.

An outbred lab culture and a set of 12 inbred lines of *M. lignano* expressing a green fluorescent protein (GFP) in all cell types, including sperm cells, allowed us to distinguish between sperm from a focal GFP-positive (hereafter denoted as GFP+) sperm donor from those of wildtype competitors with no GFP expression (hereafter GFP-). We used these inbred lines to conduct mating trials testing how variation in sperm size among the lines predicted sperm transfer success (i.e., the number of sperm stored in the partner’s sperm storage) in the presence of sperm competition. While we hypothesized an association between sperm size and sperm competitiveness, we did not have a clear prediction for the shape of this relationship as the factors determining its exact form (e.g., resource allocation among ejaculate components, sperm displacement propensity and spatial constraints for sperm storage) are largely unknown for *M. lignano*.

## Methods

### Model organism

The free-living flatworm *Macrostomum lignano* (Macrostomorpha, Platyhelminthes) is an obligatorily outcrossing simultaneous hermaphrodite found in the intertidal zones of the Northern Adriatic Sea and the Aegean Sea (Ladurner et al., 2005; Schärer et al., 2020). Laboratory cultures of *M. lignano* are kept at 20°C in glass petri dishes with artificial sea water (ASW) at a salinity of 32 ‰ and fed with the diatom *Nitzschia curvilineata*. Under laboratory conditions, *M. lignano* is found to be highly promiscuous and to mate frequently when kept in groups. Copulation in this species involves reciprocal intromission of the male copulatory organ into the female sperm-receiving and -storage organ called the antrum (Schärer et al., 2004; Vizoso et al., 2010). *M. lignano* exhibits a post-copulatory suck behaviour, in which worms bend down and place their pharynx on top of their own genital opening and appear to suck, which may have the function to remove sperm out of the antrum (Vizoso et al., 2010). The transparent body of these worms enables the observation of numerous internal structures and processes *in vivo* such as testis and ovaries, stylet morphology and sperm (both GFP+ and GFP-) stored in the female antrum (Janicke et al., 2011; Marie-Orleach et al., 2016, 2021).

The morphology of sperm in *M. lignano* is complex with two structures hypothesized to have specialized functions in the post-copulatory stages of reproduction (Visozo et al., 2010; Ladurner et al., 2005; Willems et al., 2009). The lateral bristles have been hypothesized to reduce the likelihood of sperm being removed from the antrum during a post-copulatory suck behaviour (Vizoso et al., 2010). The feeler has been observed to anchor sperm cells to the epithelium at the anterior region of the antrum (called cellular valve), where fertilization is hypothesized to occur (Vizoso et al., 2010). Given their functions, we predict selection for longer lateral bristles and feelers. Longer bristles may enable better resistance from being sucked out of the antrum and longer feelers may allow better positioning of the sperm into the cellular valve to increase the chance of fertilizing an egg. The sperm body carries a core of mitochondria and the nucleus is present in the extended sperm shaft, which also contains mitochondria. The posterior brush is made of microtubules protruding at the end of the sperm shaft. No apparent functions have been observed for sperm body, shaft and posterior brush in the female antrum so far, thus we had no clear predictions for the form of selection these sperm components. Therefore, in addition to total sperm length, we only test our predictions for feeler length and bristle length in this study.

### Experimental setup

In this study, we aimed (i) to quantify between-individual variation in sperm size and (ii) to assess the relationship between sperm size and sperm transfer success in *M. lignano*. To this end, we measured the sperm size of adult worms (N = 39) from the outbreeding BAS1 culture to quantify the natural phenotypic variation in sperm morphology in *M. lignano*. The BAS1 culture is an outbred GFP+ culture derived from backcrossing the GFP marker from a GFP+ inbred line (HUB1) onto a GFP-outbred culture for 9 generations (Marie-Orleach et al., 2016). BAS1 worms express GFP ubiquitously in all cells, including sperm cells. For our second aim, we conducted mating experiments and measured relative sperm competitiveness of worms known to vary in sperm size. Sperm size of worms cannot be measured before mating experiments, as it is a destructive process harming the worms (see next section). Hence, we used inbred lines of *M. lignano*, which are expected to show low within-but large between-line variation in sperm size. Using inbred lines also allowed us to estimate the amount of variation in sperm size attributable to genetic variance across lines.

We investigated variation in sperm size among LM lines, a set of 12 GFP+ inbred lines of *M. lignano* derived from the inbred GFP-DV lines by backcrossing the GFP marker from the HUB1 inbred line onto themselves for several generations (Marie-Orleach et al., 2017). For the sperm competition experiment, we chose 8 of the 12 LM lines with relatively narrow variation in sperm size, competing against common competitors from a single GFP-inbred line called DV1. Like other DV lines, DV1 inbred line was initiated by crossing two virgin worms taken from an outbred culture, followed by 15 generations of full-or half-sib inbreeding (For more details, refer Janicke et al., 2013 & Vellnow et al., 2017). Using worms from the GFP-DV1 line as common competitors of all LM lines allowed us to observe and distinguish GFP+sperm of LM sperm donors from the GFP-sperm of their DV1 competitors. Sperm competitiveness was estimated by calculating sperm transfer success as the proportion of GFP+ focal sperm relative to total number of sperm stored in the antrum. Sperm transfer success has been shown to positively covary with mating success and paternity share in *M. lignano* (Marie-Orleach et al., 2016), thus representing an important fitness component of the male sex function. As mating rates have been shown to be high (Schärer et al., 2004), using unmated (virgin) worms would not represent a realistic scenario to study reproductive traits in this species. Therefore, worms were raised in small groups of 3 worms before the mating trials so that they were sexually experienced and had reached a biologically realistic steady state of sperm and egg reserves ready to be donated and fertilized, respectively.

### Sperm size measurements

Measurements taken from each sperm cell included the total sperm length, measured from anterior tip of feeler to the posterior tip of brush (Figure 1), and the length of anterior feeler and lateral bristles (mean length of the two bristles). Measurements were done using an established technique previously used in *M. lignano* (Janicke and Schärer, 2010). Adult worms were isolated for a day in wells of a 24-well plate to allow them to restore sperm reserves in their seminal vesicles, where they are stored until the sperm is transferred to a partner during copulation. The tail plates of the worms consisting the seminal vesicle were amputated and transferred onto a microscopic slide with 1 μl of ASW medium, rupturing the seminal vesicle and causing sperm to spill out into the surrounding medium. This restricted the movement of sperm cells greatly, facilitating the imaging of sperm morphology with a digital camera (The Imaging Source, DFK 41BF02) connected to a light microscope (Leica DM2500, Leica Micro-systems, Germany) and imaging software BTV Pro 6.0b7 (http://www.bensoftware.com/).

**Figure 1.**
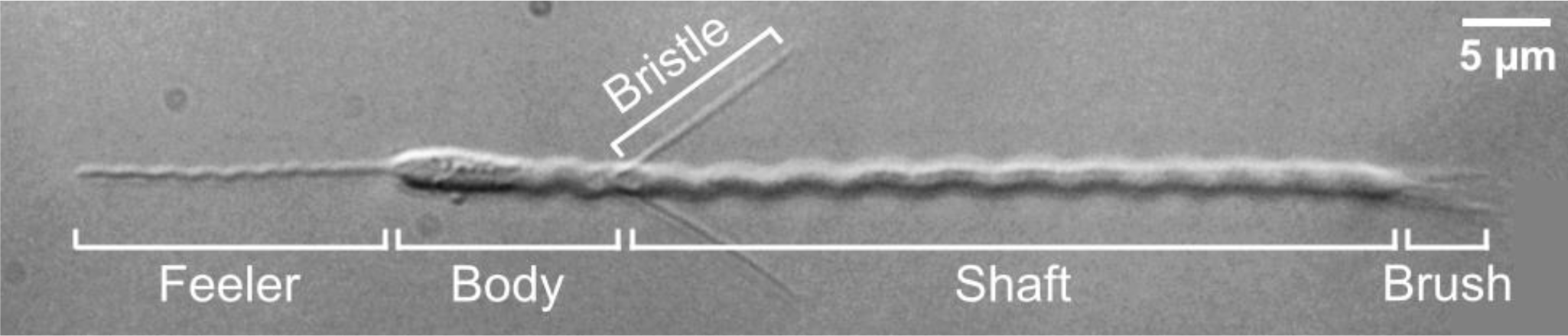
Morphology of a mature sperm cell in *Macrostomum lignano*.

We took digital micrographs of sperm cells at 1000x magnification and the micrographs were analyzed in ImageJ 1.53c using the ObjectJ plugin (https://sils.fnwi.uva.nl/bcb/objectj/). About 15 sperm cells were measured from all individuals that were measured as part of this study. This number has been shown to be adequate to provide an accurate estimate of the individual mean values of sperm length (Janicke and Schärer, 2010). Additionally, we estimated the measurement error in this method by calculating the repeatability when the total sperm length, bristle length and feeler length of the same sperm cell was measured thrice for a total of 10 randomly chosen sperm cells each from different worms.

### Rearing conditions

On day 1, 200 GFP+ adults from each of the 12 LM lines and 2400 GFP-adults of the DV1 line were distributed into petri dishes, 100 adults in each, for egg laying with 20 ml ASW and ad libitum algae. On day 3, we removed all adults from the dishes controlling the age differences in the resulting juveniles to less than 48 hours. On day 9, we sampled the hatchlings to form rearing groups of three hatchlings, all from the same inbred line. In total, we formed 30 such groups for each LM line and 360 groups from the DV1 line. The groups were placed in wells of 24-well plates with 1.5 ml of 32‰ ASW and ad libitum algae. To control for any environmental effects during development, every 24-well plate contained 2 groups from each of the LM lines in randomly assigned wells. All groups were transferred to fresh medium and algae every 6-10 days until adulthood. During this period, sperm size of the 12 LM lines were measured from a different set of adult worms (56 - 63 days old) and 8 lines with relatively narrow within-line variation in sperm morphology were chosen for the sperm competition experiment.

### Sperm competition experiment

On days 35 through 55, we assessed sperm transfer success of worms from the 8 LM lines when competing against the common competitor DV1 line. On every experiment day, we formed ‘mating groups’ by moving one randomly picked adult from a LM rearing group, verified for GFP expression, to a fresh well along with three DV1 worms. Thus, a mating group consisted of one GFP+ focal worm from one of the 8 LM lines and three GFP-worms of a DV1 rearing group, and each focal LM worm competed against two DV1 competitors for access to eggs of a third DV1 worm. On the following day, the focal LM worm was removed from the experimental group.

Immediately after the 24-hour mating trial, the number of GFP- and GFP+ sperm in the female antrum of the three DV1 partners was assessed, as previously reported (Janicke et al., 2013). We used a digital video camera (ORCA-Flash 4.0 V2: C11440-22CU, Hamamatsu corporation) connected to a Leica DM2500 microscope (Leica Micro-systems, Germany) and the software Leica Applications Suite X (LAS X, Leica Micro-systems, Germany) to make movies of the female antrum. For each DV1 partner, we recorded a movie of the female antrum by focusing through the antrum under differential interference contrast illumination to be able to see all the sperm in the antrum (movie 1). Immediately after, we recorded a second movie of the female antrum under epifluorescence illumination to visualize the GFP+ sperm that were transferred and stored by the focal worm (movie 2). We analysed the movies using VLC media player and counted, for each focal donor, the total number of sperm (from movie 1) and the number of GFP+ sperm (from movie 2) from the focal donor in the female antrum of all its potential partners.

Similar to previous studies with *M. lignano*, we encountered worms with an egg (or eggs) in the antrum (i.e., 263 out of 720 DV1 worms used in the experiment). In the presence of eggs in the antrum, it is not possible to reliably count all sperm that remain in the antrum under interference contrast illumination, which meant we could only count the GFP+ sperm in these partners. Previous studies extrapolated the total number of sperm in such worms by using the average value of the total number of sperm in all the other worms (Marie-Orleach at al., 2016, 2021). In this study, the number of GFP+ sperm was sometimes greater than the total number of sperm resulting in proportion values higher than 1. Therefore, we performed all analysis excluding worms that had eggs in their female antrum.

### Statistical analysis

All statistical analysis for this study were performed in R (Version 4.2.3). Variation in sperm size The total natural variation in sperm length in *M. lignano* was assessed by calculating the coefficient of variation for each sperm morphological trait in the outbred BAS1 culture. We tested for between-individual differences on all sperm traits using Kruskal Wallis rank sum tests with individual identifier as the predictor variable.

Repeatabilities for the measurement method were calculated for 3 measurements of each sperm cell with the rptGaussian function in the *rptR* package (V0.9.22, Stoffel et al., 2017), using 1000 parametric bootstraps to evaluate the uncertainty in the repeatability estimate. All further analysis were done using the arithmetic mean of sperm measurements obtained from each individual. We estimated the proportion of phenotypic variation in sperm size that is explained by the LM line identity, i.e., variance that can be attributed to the genotype of the male, using a linear mixed models with LM line identifier and individual worm identifier as random effects and trait length as dependent variable. The linear mixed models were built using the ‘lmer’ function in the *lme4* package in R (V1.1-32, Bates et al., 2015), assuming the individual worm identifier to be nested within the LM line identifier.

### Relationship between sperm length and sperm transfer success

Initially, we intended to run 240 sperm competition trials (30 per line), but we lost 4 replicates due to handling errors. Moreover, 263 (36.5%) of the 720 DV1 partners had one or more eggs in their antrum, resulting in the loss of 24 replicates where all partners had eggs in their antrum at the time of observation. Previous studies have reported comparable frequencies for the occurrence of eggs in the female antrum in *M. lignano* (Marie-Orleach at al., 2016, 2021). Thus, the final analysis was performed with a total of 212 replicates (24 to 29 replicates per line, mean = 26.5). For each replicate, we summed the total number of sperm and the number of focal GFP+ sperm found in the three partners (only 2 or 1 in cases of eggs or worms were lost). Sperm transfer success of a GFP+ focal worm was calculated as the proportion of GFP+ sperm among the total number of sperm stored in the antrum of all the partners.

Firstly, we assessed the differences in sperm transfer success among the LM lines. We fitted binomial Generalized linear models (GLM) using the *lme4* package (V1.1-32, Bates et al., 2015), with line identity as a categorical fixed effect and the relative number of GFP+ and GFP-sperm with the ‘cbind’ function as the response variable representing sperm transfer success. We observed over-dispersion in the binomial GLM (dispersion ratio = 9.344), which may arise from the variation in the total number of sperm observed in partners’ antrum (mean ± standard deviation = 44 ± 24). Therefore, we refitted the models assuming a quasibinomial data distribution. Additionally, we tested if the total number of sperm observed in partners’ antrum, sampled to estimate sperm transfer success differed between the LM lines, as this may suggest that systematic difference in sperm production between mating groups of different LM lines. For this, we built a generalized linear model with total number of sperm sampled as the dependent variable and the LM line identifier as the categorical predictor variable, assuming a quasipoisson data distribution to account for over-dispersion (dispersion parameter = 13.42).

Secondly, we assessed the relationship between sperm transfer success and total sperm length. Given their functional significance in post-copulatory sexual selection, we performed the assessment also for lateral bristle lengths and anterior feeler lengths. As mentioned earlier, sperm length measurements are invasive requiring the amputation of the tail plate so that we could not measure sperm morphology of focal worms used in the mating experiments. Therefore, we used the mean value of each line (hereafter, called “line means”) as our measure of trait length. This meant that all focal worms from the same LM line were assumed to express similar length, for all three traits. However, to account for the variation within lines such that mean values of lines with higher precision are given relatively higher weight compared to those with lower precision, we used the inverse of the standard deviation in sperm length within lines as a weighing factor in our models.

We built two quasibinomial regression models, one first-order linear and one including, in addition, a second-order quadratic term, with individual sperm transfer success as the response variable and line means of sperm traits as a continuous predictor variable. In order to assess the potential for a hump-shaped relationship between sperm length and sperm transfer success, we used a quadratic polynomial of line means as the independent predictor variable. We performed a likelihood ratio test to infer whether adding the quadratic term provides a better model fit. In addition, we compared the goodness of fit of the two models by using the Akaike information criterion accounting for the over-dispersion in the data (quasi-AIC), as implemented in the *bbmle* package (V1.0.25, Bolker, 2022) in R.

## Results

The measurement repeatabilities were high for all three sperm traits measured (Table S1). The total and bristle length were highly repeatable whereas repeatability was lower for the feeler length (Table S1). This was expected as undulations of the feelers caused difficulties in accurate measurement of the length.

### Variation in sperm size among individuals and inbred lines

In the outbred BAS1 culture, we observed large variation in sperm length ranging from 70 to 90µm, approximately. We found significant between-individual variation in all sperm traits measured (Table S1, also see Figure S1 for variation in total sperm length). The coefficients of variation (CV) of all sperm traits were comparable (4.31 to 6.10 %). The variation in sperm size within individuals (WCV) was on average lower than the overall variation observed in the culture (Table S1).

As expected, the combined variation in sperm size observed in the LM lines approximately covered the overall phenotypic variation measured from the outbred culture (Figure 2). The sperm size of the DV1 line (i.e. used as common competitors in the mating trials) corresponded to the average sperm size found in LM lines (Figure 2). Therefore, LM lines had larger, similar and smaller sperm size to DV1, allowing us to study the potential role of competitor’s sperm size on sperm transfer success. We found a significant between-line variation in all sperm traits measured (total sperm length: Kruskal-wallis test, χ^2^ = 65.34, df = 11, *P* = 9.3e^-10^). The linear mixed models revealed that a substantial portion of phenotypic variation in all sperm traits could be explained by LM line identity (Table S2), explaining 45 % of variation in total sperm length.

**Figure 2.**
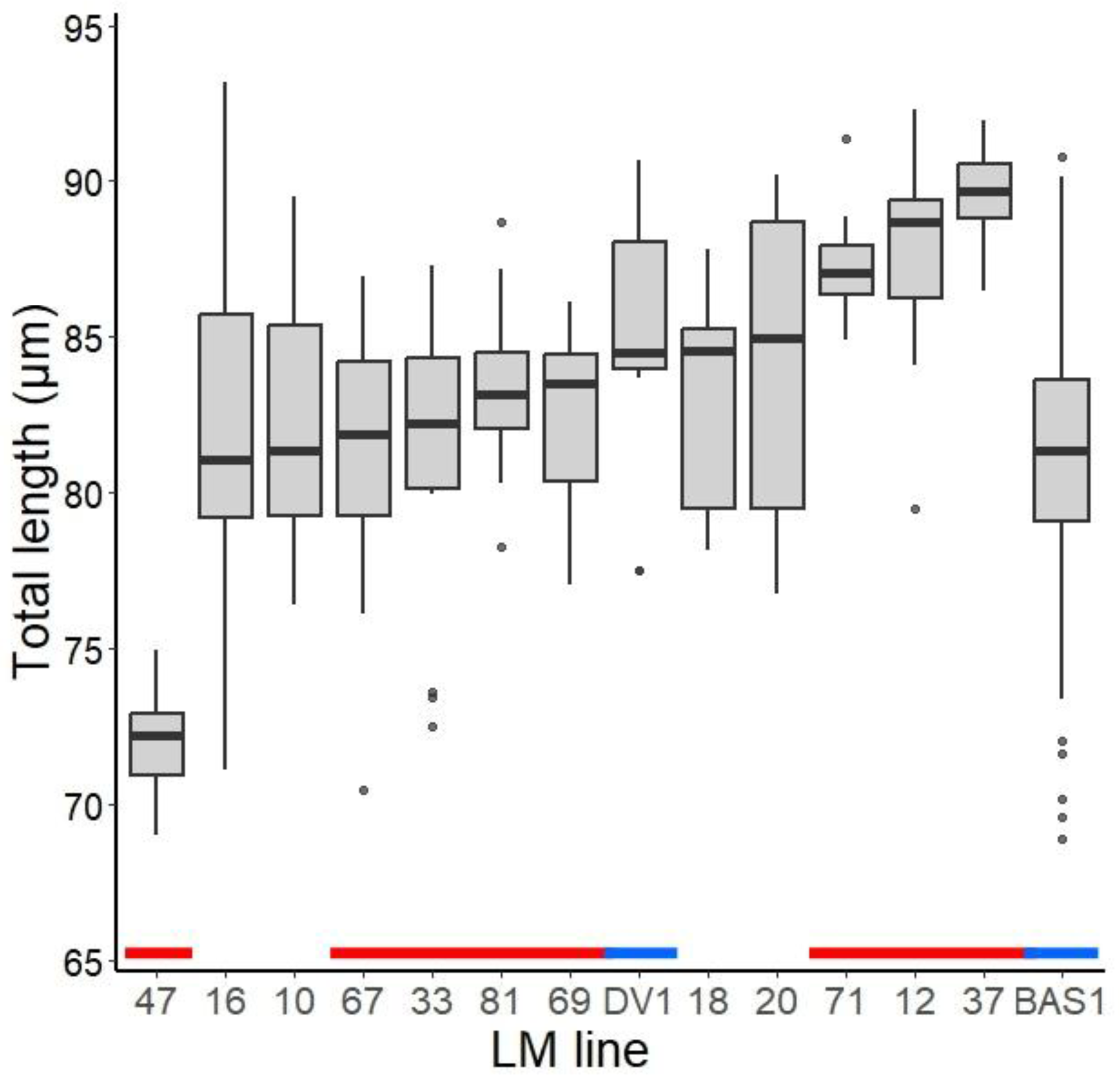
Variation in total sperm length among the 12 LM lines, DV1 line and the outbred BAS1 culture. LM lines are arranged in increasing order of median total sperm length. Red bars indicate the 8 LM lines chosen for the sperm competition experiment and the blue bars indicate the DV1 and BAS sperm size.14-15 worms were measured for each line and 39 worms for the BAS1 culture. About 15 sperm cells were measured for each worm in both BAS1 and LM lines.

### Sperm size and sperm transfer success

The generalized linear model testing differences between mating groups of LM lines revealed that the total number of sperm sampled did not differ between the mating groups of LM lines (χ^2^ = 7.30, df = 7, *P* = 0.40), showing that our measures of sperm transfer were not biased by the number of sperm sampled. We observed a significant between-line variation in sperm transfer success of LM focal worms (line identifier: χ^2^ = 52.07, df = 7, *P* = 5.65e^-9^). We detected significant negative relationships in the first-order linear models between sperm transfer success and all three sperm traits modelled (Table 1). The second-order quadratic models revealed a significant non-linear relationship with intermediate lengths having higher sperm transfer success for all three sperm traits (Table 1, Figure 3 and S2). Additionally, likelihood-ratio tests suggest that the quadratic regression models provided significantly better fits to the data compared to the linear regression models (Table 1).

**Figure 3.**
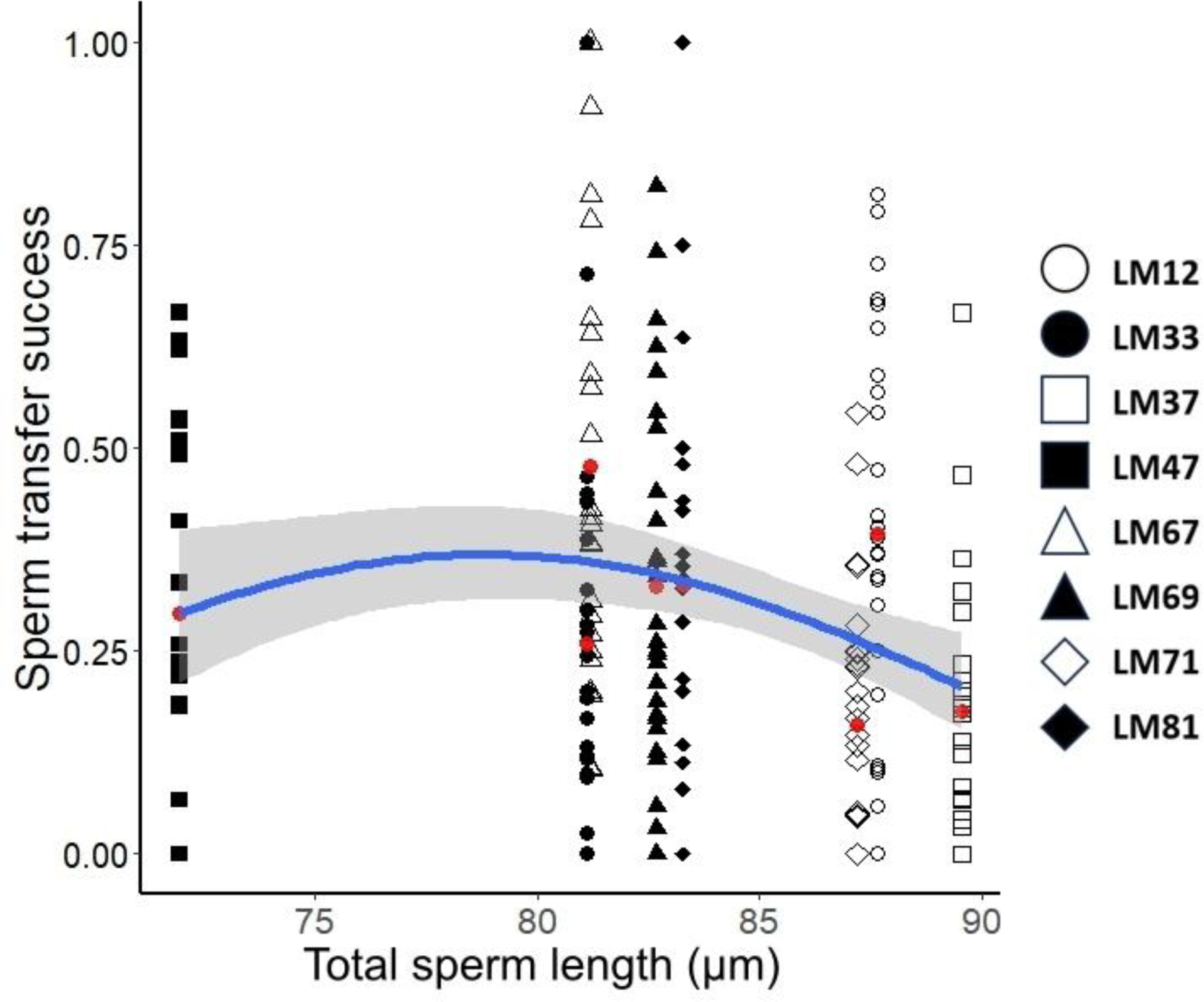
Relationship between total sperm length and sperm transfer success. Symbols indicate different LM lines (see legend). Red dots indicate mean sperm transfer success in each line. Solid blue line shows the fit of the quadratic model using all data points, and grey regions indicate the 95% confidence intervals.

**Table 1.** Statistical analysis of the relationship between sperm traits and sperm transfer success. Results of ANOVA Wald tests for linear and quadratic polynomial regressions are shown together with likelihood ratio tests comparing the two models for all sperm morphological traits. Significant *P* values are shown in bold.

| *Sperm trait* | Linear model |  |  |  | Quadratic model |  |  |  | LRT |  |
| --- | --- | --- | --- | --- | --- | --- | --- | --- | --- | --- |
| | $\chi^2$ | *P* | R<sup>2</sup> | qAIC | $\chi^2$ | *P* | R<sup>2</sup> | qAIC | $\chi^2$ | *P* |
| **Total sperm length** | 9.497 | **0.002** | 0.039 | 300.88 | 7.859 | **0.005** | 0.073 | 294.45 | 37.10 | **0.003** |
| **Total sperm length<sup>2</sup>** |  |  |  |  | 8.415 | **0.004** |  |  |  |  |
| **Feeler length** | 8.903 | **0.003** | 0.037 | 291.77 | 3.560 | 0.059 | 0.054 | 289.72 | 55.33 | **0.043** |
| **Feeler length<sup>2</sup>** |  |  |  |  | 4.010 | **0.045** |  |  |  |  |
| **Bristle length** | 4.866 | **0.027** | 0.020 | 300.99 | 6.547 | **0.011** | 0.049 | 296.01 | 170.00 | **0.008** |
| **Bristle length<sup>2</sup>** |  |  |  |  | 6.790 | **0.009** |  |  |  |  |

## Discussion

Sperm size has repeatedly been found to evolve in response to sperm competition across species (Balshine et al., 2001; Godwin et al., 2017; LaMunyon and Ward, 2002; Tourmente et al., 2011). However, our understanding of how sperm size affects sperm competitiveness and hence male reproductive performance within species is limited. In this study, we observed a hump-shaped relationship between post-mating success and sperm size with sperm of intermediate size having on average the highest competitiveness in the simultaneously hermaphroditic flatworm *M. lignano*. Our results provide support for the hypothesis that sperm competition may favour the evolution of intermediate sperm size (Calhim et al., 2007; Parker, 1993). In the following, we discuss our findings and point to the strengths and limitations of our study.

### Sperm transfer success and sperm size

Non-linear selection towards an optimal sperm size has been suggested mainly by comparative studies (Kleven et al., 2008; Lüpold et al., 2009), inferred from a reduction in sperm size variation in species with higher levels of sperm competition. Lymbery et al. (2018) observed non-linear selection with an intermediate peak on canonical axes of multiple sperm traits including sperm length, swimming velocity and swimming path linearity in the mussel, *Mytilus galloprovincialis*. In *M. lignano*, a recent study tested whether various sperm traits predict male reproductive performance (Marie-Orleach et al., 2023) and found no significant non-linear relationship between sperm morphology and sperm transfer efficiency - a fitness component comparable to sperm transfer success assessed in this study. This previous study did not explore the importance of total sperm length and studied purely phenotypic effects of sperm morphological traits on male fitness components, which does not allow inference of the evolutionary trajectories. In the present study, we aimed at filling this gap by exploring how genetic variance in total sperm length and other traits relates to sperm transfer success among inbred LM lines (discussed further below). Estimates of mean sperm length obtained from inbred lines correspond to breeding values, which allowed us to investigate how sperm competition selects on standing genetic variance in sperm length. Our approach demonstrates that selection arising from sperm transfer success favours the evolution towards intermediate length for all three sperm traits most relevant to post-copulatory sexual selection.

In our study, we focused on sperm transfer success to quantify potential selection on sperm size arising from competition to deposit sperm in the female reproductive tract, which is a crucial step in order to obtain reproductive success through male function. In *M. lignano*, sperm transfer efficiency (i.e., sperm transfer success per successful mating) has been estimated to explain 27% of the variation in male reproductive success (Marie-Orleach et al., 2021). The fitness component that explained, with 37%, most variation in male reproductive success was sperm fertilizing efficiency (i.e., the proportion of sired offspring relative to the number of sperm stored in the female sperm storage organ). However, sperm transfer success has been shown to strongly predict the proportion of sired offspring in *M. lignano* (Marie-Orleach et al., 2016) indicating that sperm transfer success is a good proxy for post-mating performance in the species. Nevertheless, we acknowledge that other aspects, such as sperm fertilizing efficiency, may also affect selection on sperm traits and remain unexplored in our study.

All three sperm traits exhibited significant negative linear relationships with sperm transfer success. Such negative associations have been suggested to support the theory that sperm competition generates selection for increased number of sperm, which trades-off with sperm size resulting in smaller sperm (Gage and Morrow, 2003; Garcìa-González and Simmons, 2007). However, hump-shaped relationships provided the best fit to sperm transfer success for total length, bristle length and feeler length, suggesting selection for intermediate sperm size. Selection for increased sperm number alone cannot explain this result satisfactorily. Selection for intermediate sperm size, however, was predicted by theoretical models when fertilization efficiency of sperm increased with a decreasing gradient in relation to size (Parker 1993, Parker and Begon, 1993, Parker et al., 2010). These models assumed that larger sperm have a fertilization advantage over smaller sperm but increase in mass of each sperm traded off against other male reproductive expenditures, such as sperm number and mate acquisition. We suspect that the lower success of individuals with particularly long sperm observed in our study may be explained by this trade-off between sperm size and number and that selection for intermediate sperm size may provide a balance between the selective forces for more competitive, but fewer, large sperm and numerous, but less competitive, small sperm.

The selection for intermediate sperm size could also arise from the female environment where sperm compete, as sperm size of DV1 fell in the intermediate range of the LM sperm size distribution. Sperm size has been shown to covary with female reproductive tract (or its components) across diverse taxa (Minder et al., 2005; Morrow and Gage, 2000; Pitnick et al., 1999, 2003; Briskie and Montgomerie 1992) and to evolve in response to selection on the female reproductive tract in *Drosophila melanogaster* suggesting a role of female sperm choice for shaping sperm morphology (Miller and Pitnick, 2002). Cryptic female choice by DV1 recipients in terms of a choice for sperm size that were similar to their own may have favoured intermediate sperm size in our experiment. However, the female antrum in *M. lignano* is a simple organ with a single chamber to receive and store sperm. The cell membrane along the wall of the antrum is convoluted and highly flexible, which is required for the passage of eggs (Vizoso et al., 2010). Given the simple structure and flexibility in size, we do not have a clear mechanistic explanation for how the antrum may exert female choice on sperm in relation to size.

Different components of sperm morphology may have very different consequences for sperm competitiveness, such that different traits show divergent associations to male reproductive success within species. For example, in the red deer (*Cervus elaphus*), sperm head and tail length were positively associated with sperm swimming velocity and male fertility, but mid-piece length showed a negative correlation with male fertility (Malo et al., 2006). In *M. lignano*, bristle and feeler length have been hypothesized to play active roles in post-copulatory stages, preventing removal of sperm from the female antrum and enabling anchoring of sperm to the cellular valve, respectively (Vizoso et al., 2010). Selection arising from sperm competition can be expected to be stronger on these two traits compared to others and we predicted linear selection for longer bristles and feelers. Our results showed non-linear selection for intermediate length of these traits. This finding might be the consequence of strong positive correlations between the length of both traits and total sperm length (Janicke et al., 2010), for which we also found support for stabilising selection.

Using inbred worms provided us with groups of individuals that are expected to be genetically similar and we expected them to have a relatively narrow range of sperm size within lines and significant differences among lines. While we found clear differences among lines, we also observed substantial variation in sperm length within lines. This might explain the high variation in sperm transfer success within lines, reiterating the significance of stochastic and non-genetic sources of variation in the measurements of post-mating success (Marie-Orleach et al., 2021). Moreover, *M. lignano* exhibits a high mating rate, such that the focal worms are expected to have copulated repeatedly with each mating partner in the 24-hour mating trials. Our experimental setup did not allow to control the number of copulations and we acknowledge that variation in sperm transfer success may also represent varying levels of multiple mating.

The inbred lines used in the study were derived from four geographically distinct populations of *M. lignano* (Vellnow et al., 2017) and are expected to differ genetically, resulting in variation between the LM lines. Therefore, phenotypic differences other than sperm size arising from focal LM genotype and/or its interaction with the partner DV1 genotype, could have contributed to between-line variation in sperm transfer success in the LM lines. Moreover, partner genotypes have been shown to affect reproductive morphology and behaviour in *M. lignano* (Marie-Orleach et al., 2017) but evidence for the influence of partner genotype on fitness components, such as mating success and sperm transfer success, is still lacking. Thus, as true for most studies on selection gradients, our results do not allow to establish a causal link between sperm size and sperm transfer success.

A shortcoming of our study was a discontinuous gradient in sperm size among LM lines with a particularly big difference between the line with the shortest (LM47) and the line with the second shortest sperm (LM67). This resulted in the LM 47 being the only representative of the lower end of the spectrum in sperm size. However, we decided not to remove LM47 from the analysis because our aim was to cover the largest possible variation in sperm size that could be observed in the outbred BAS1 culture of *M. lignano*, which itself is likely to harbour only a fraction of the variation in sperm size compared to natural populations. Another potential shortcoming of our study is that the variation in sperm length within inbred lines was unexpectedly high. We did not measure the actual sperm size of the focal LM worms used in the mating experiments. Given that focal worms were taken from inbred lines with relatively narrow range of variation, we expect the actual sperm size of the focal worms would not differ substantially from the values used in our analysis. We also accounted for among-line differences in the variance of sperm length by adding the inverse of standard deviations in sperm size as a weighting factor in our model. Our results remained qualitatively unaffected without the weighting factor in the model.

### Intra-specific variation in sperm size

Intra-specific variation in sperm size is a frequently observed phenomenon across animal taxa (reviewed in Pitnick et al., 2009). We also observed substantial variation in sperm size in both BAS1 culture and among the LM lines. The inbred line identity explained a substantial fraction of sperm trait variation (45% variation in total sperm length). This portion of variation is expected to be attributable to the genetic variance between the tested LM lines and is an estimate of the total genetic variance (equivalent to the broad-sense heritability) in *M. lignano*. This result is in line with the commonly observed high heritability in sperm size across animals (Simmons and Moore, 2009). In addition, we also observed substantial variation in sperm size within LM lines, with some lines expressing a range of sperm length equivalent to that of BAS1. The unusual genetic system of *M. lignano* might contribute to this relatively large within-line variation with karyotype polymorphisms frequently observed. Worms of this species have been reported to carry 2n = 8, 2n = 9 or 2n = 10 chromosomes with rare cases of other karyotypes (Zadesenets et al., 2016, 2020). The BAS1 culture has been generated such that all worms carry the normal karyotype of 2n=8 (Vellnow et al., 2018). However, the karyotypes of the LM lines have not yet been explored. Ancestral DV lines were derived from wild populations and karyotype polymorphisms may still persist in the LM lines, and thus contribute to the observed phenotypic variation. Karyotype polymorphism could result in increased variation in mRNA dosages of genes influencing sperm morphology and additional copies of chromosomes may account for more genetic material that has to be physically packed within the sperm shaft.

## Conclusion

We used *in vivo* sperm counting to assess how genetic variance in sperm size translates into post-mating success in an outcrossing simultaneous hermaphrodite. Our finding of a hump-shaped association between sperm size and post-mating success suggests that sperm competition may favour an intermediate sperm size. We speculate that such stabilizing selection most likely results from a trade-off between sperm size and sperm number in *M. lignano*, as predicted by theory. Our study is one among the few to explore and present evidence for non-linear selection arising from post-copulatory sexual selection on sperm size and calls for future studies to consider divergent forms of selection on ejaculate traits.

## Acknowledgements

We thank Lukas Schärer for significant contributions to conceiving the study and designing the experiment. We also thank Gudrun Viktorin, Jurgen Hottinger and Daniel Lüscher for technical support, and Yasmin Picton for administrative support.

## References

Balshine, S., Leach, B. J., Neat, F., Werner, N. Y., and Montgomerie, R. (2001). Sperm size of African cichlids in relation to sperm competition. Behavioral Ecology, 12(6), 726–731.

Bates, D., Maechler, M., Bolker, B., and Walker, S. (2015). Fitting Linear Mixed-Effects Models Using lme4. Journal of Statistical Software, 67(1), 1–48. doi:10.18637/jss.v067.i01.

Bennison, C., Hemmings, N., Slate, J., and Birkhead, T. (2015). Long sperm fertilize more eggs in a bird. Proceedings of the Royal Society B: Biological Sciences, 282(1799), 20141897.

Birkhead, T., and Moller, A. P. (1999). Sperm Competition and Sexual Selection. Academic Press, London.

Bolker, B, R Development Core Team (2022). bbmle: Tools for General Maximum Likelihood Estimation_. R package version 1.0.25, <https://CRAN.R-project.org/package=bbmle>.

Briskie, J. V., & Montgomerie, R. (1992). Sperm size and sperm competition in birds. Proceedings of the Royal Society of London. Series B: Biological Sciences, 247(1319), 89–95.

Briskie, J. V, Montgomerie, R., and Birkhead, T. (1997). The Evolution of Sperm Size in Birds. Evolution, 51(3), 937–945.

Calhim, S., Immler, S., and Birkhead, T. (2007). Postcopulatory Sexual Selection Is Associated with Reduced Variation in Sperm Morphology. PLoS ONE, 2(5), e413.

Cascio Sætre, C. Lo, Johnsen, A., Stensrud, E., and Cramer, E. R. A. (2018). Sperm morphology, sperm motility and paternity success in the bluethroat (*Luscinia svecica*). PLoS ONE, 13(3), e0192644.

Collet, J., Richardson, D. S., Worley, K., and Pizzari, T. (2012). Sexual selection and the differential effect of polyandry. Proceedings of the National Academy of Sciences, 109(22), 8641–8645.

Cramer, E. R., Laskemoen, T., Kleven, O., LaBarbera, K., Lovette, I. J., and Lifjeld, J. T. (2013). No evidence that sperm morphology predicts paternity success in wild house wrens. Behavioral Ecology and Sociobiology, 67, 1845–1853.

Fitzpatrick, John L., and Stefan Lüpold. (2014) Sexual selection and the evolution of sperm quality. Molecular Human Reproduction, 20(12), 1180–1189.

Gage, M. J. G., and Freckleton, R. P. (2003). Relative testis size and sperm morphometry across mammals: No evidence for an association between sperm competition and sperm length. Proceedings of the Royal Society B: Biological Sciences, 270(1515), 625–632.

Gage, M. J., and Morrow, E. H. (2003). Experimental evidence for the evolution of numerous, tiny sperm via sperm competition. Current Biology, 13(9), 754–757.

García-González, F., and Simmons, L. W. (2007). Shorter sperm confer higher competitive fertilization success. Evolution, 61(4), 816–824.

Godwin, J. L., Vasudeva, R., Michalczyk, Ł., Martin, O. Y., Lumley, A. J., Chapman, T., and Gage, M. J. (2017). Experimental evolution reveals that sperm competition intensity selects for longer, more costly sperm. Evolution Letters, 1(2), 102–113.

Immler, S., and Birkhead, T. R. (2007). Sperm competition and sperm midpiece size: no consistent pattern in passerine birds. Proceedings of the Royal Society B: Biological Sciences, 274(1609), 561–568.

Immler, S., Calhim, S., and Birkhead, T. R. (2008). Increased postcopulatory sexual selection reduces the intramale variation in sperm design. Evolution, 62(6), 1538–1543.

Janicke, T., and Schärer, L. (2010). Sperm competition affects sex allocation but not sperm morphology in a flatworm. Behavioral Ecology and Sociobiology, 64, 1367–1375.

Janicke, T., Sandner, P., and Schärer, L. (2011). Determinants of female fecundity in a simultaneous hermaphrodite: the role of polyandry and food availability. Evolutionary ecology, 25, 203–218.

Janicke, T., Marie-Orleach, L., De Mulder, K., Berezikov, E., Ladurner, P., Vizoso, D. B., and Schärer, L. (2013). Sex allocation adjustment to mating group size in a simultaneous hermaphrodite. Evolution, 67(11), 3233–3242.

Janicke, T., Marie-Orleach, L., De Mulder, K., Berezikov, E., Ladurner, P., Vizoso, D. B., & Schärer, L. (2013). Sex allocation adjustment to mating group size in a simultaneous hermaphrodite. Evolution, 67(11), 3233–3242.

Kahrl, A. F., Kustra, M. C., Reedy, A. M., Bhave, R. S., Seears, H. A., Warner, D. A., and Cox, R. M. (2021). Selection on sperm count, but not on sperm morphology or velocity, in a wild population of anolis lizards. Cells, 10(9), 2369.

Kleven, O., Laskemoen, T., Fossøy, F., Robertson, R. J., and Lifjeld, J. T. (2008). Intraspecific variation in sperm length is negatively related to sperm competition in passerine birds. Evolution, 62(2), 494–499.

Ladurner, P., Schärer, L., Salvenmoser, W., and Rieger, R. M. (2005). A new model organism among the lower Bilateria and the use of digital microscopy in taxonomy of meiobenthic Platyhelminthes: *Macrostomum lignano*, n. sp. (Rhabditophora, Macrostomorpha). Journal of Zoological Systematics and Evolutionary Research, 43(2), 114–126.

LaMunyon, C. W., and Ward, S. (2002). Evolution of larger sperm in response to experimentally increased sperm competition in *Caenorhabditis elegans*. Proceedings of the Royal Society of London. Series B: Biological Sciences, 269(1496), 1125–1128.

Lenth R. (2023). _emmeans: Estimated Marginal Means, aka Least-Squares Means_. R package version 1.8.6, <https://CRAN.R-project.org/package=emmeans>.

Lüpold, S., Linz, G. M., and Birkhead, T. R. (2009). Sperm design and variation in the New World blackbirds (Icteridae). Behavioral Ecology and Sociobiology, 63, 899–909.

Lüpold, S., Pitnick, S., Berben, K. S., Blengini, C. S., Belote, J. M., and Manier, M. K. (2013). Female mediation of competitive fertilization success in Drosophila melanogaster. Proceedings of the National Academy of Sciences, 110(26), 10693–10698.

Lymbery, R. A., Kennington, W. J., & Evans, J. P. (2018). Multivariate sexual selection on ejaculate traits under sperm competition. The American Naturalist, 192(1), 94–104.

Malo, A. F., Gomendio, M., Garde, J., Lang-lenton, B., Soler, A. J., and Roldan, E. R. S. (2006). Sperm design and sperm function. Biology Letters, 2(2), 246– 249.

Marie-Orleach, L., Janicke, T., and Schärer, L. (2013). Effects of mating status on copulatory and postcopulatory behaviour in a simultaneous hermaphrodite. Anim. Behav. 85:453–461.

Marie-Orleach, L., Janicke, T., Vizoso, D. B., David, P., and Schärer, L. (2016). Quantifying episodes of sexual selection: Insights from a transparent worm with fluorescent sperm. Evolution, 70(2), 314–328.

Marie-Orleach, L., Vogt-Burri, N., Mouginot, P., Schlatter, A., Vizoso, D. B., Bailey, N. W., and Schärer, L. (2017). Indirect genetic effects and sexual conflicts: Partner genotype influences multiple morphological and behavioral reproductive traits in a flatworm. Evolution, 71(5), 1232–1245.

Marie-Orleach, L., Vellnow, N., and Schärer, L. (2021). The repeatable opportunity for selection differs between pre-and postcopulatory fitness components. Evolution Letters, 5(1), 101–114.

Marie-Orleach, L., Hall, M. D., and Schärer, L. (2023). Contrasting the form and strength of pre-and postcopulatory sexual selection in a transparent worm with fluorescent sperm. Evolution - International Journal of Organic Evolution (In press).

Miller, G. T., and Pitnick, S. (2002). Sperm-Female Coevolution in Drosophila. Science, 298(5596), 1230–1233.

Minder, A. M., Hosken, D. J., and Ward, P. I. (2005). Co-evolution of male and female reproductive characters across the Scathophagidae (Diptera). Journal of Evolutionary Biology, 18(1), 60–69.

Morrow, E. H., and Gage, M. J. G. (2000). The evolution of sperm length in moths. Proceedings of the Royal Society B: Biological Sciences, 267(1440), 307–313.

Morrow, E. H., and Gage, M. J. G. (2001). Sperm competition experiments between lines of crickets producing different sperm lengths. Proceedings of the Royal Society B: Biological Sciences, 268(1482), 2281–2286.

Parker, G. A. (1970). Sperm competition and its evolutionary consequences in the insects. Biological reviews, 45(4), 525–567.

Parker, G. A. (1982). Why are there so many tiny sperm? Sperm competition and the maintenance of two sexes. Journal of Theoretical Biology, 96(2), 281–294.

Parker, G. A. (1993). Sperm competition games: Sperm size and sperm number under adult control. Proceedings of the Royal Society B: Biological Sciences, 253(1338), 245–254.

Parker, G. A., and Begon, M. E. (1993). Sperm competition games: sperm size and number under gametic control. Proceedings of the Royal Society of London. Series B: Biological Sciences, 253(1338), 255–262.

Parker, G. A., Immler, S., Pitnick, S., and Birkhead, T. (2010). Sperm competition games: Sperm size (mass) and number under raffle and displacement, and the evolution of P2. Journal of Theoretical Biology, 264(3), 1003–1023.

Pélissié, B., Jarne, P., Sarda, V., and David, P. (2014). Disentangling precopulatory and postcopulatory sexual selection in polyandrous species. Evolution, 68(5), 1320–1331.

Pitnick, S., Hosken, D. J., and Birkhead, T. (2009). Sperm morphological diversity, In Sperm biology (pp. 69-149). Academic Press, London.

Pitnick, S., Markow, T., and Spicer, G. S. (1999). Evolution of Multiple Kinds of Female Sperm-Storage Organs in Drosophila. Evolution, 53(6), 1804–1822.

Pitnick, S., Miller, G. T., Schneider, K., and Markow, T. A. (2003). Ejaculate-female coevolution in *Drosophila mojavensis*. Proceedings of the Royal Society B: Biological Sciences, 270(1523), 1507–1512.

Pitnick, S., Wolfner, M. F., and Suarez, S. S. (2009). Ejaculate–female and sperm– female interactions. In Sperm biology (pp. 247–304). Academic Press, London.

Pizzari, T., and Parker, G. A. (2009). Sperm competition and sperm phenotype. In Sperm biology (pp. 207–245). Academic Press, London.

Radwan, J. (1996). Intraspecific variation in sperm competition success in the bulb mite: a role for sperm size. Proceedings of the Royal Society of London. Series B: Biological Sciences, 263(1372), 855–859.

Rowe, M., Van Oort, A., Brouwer, L., Lifjeld, J. T., Webster, M. S., Welklin, J. F., and Baldassarre, D. T. (2022). Sperm numbers as a paternity guard in a wild bird. Cells, 11(2), 231.

Schärer, L., Brand, J. N., Singh, P., Zadesenets, K. S., Stelzer, C. P., and Viktorin, G. (2020). A phylogenetically informed search for an alternative Macrostomum model species, with notes on taxonomy, mating behavior, karyology, and genome size. Journal of Zoological Systematics and Evolutionary Research, 58(1), 41–65.

Schärer, L., Joss, G., and Sandner, P. (2004). Mating behaviour of the marine turbellarian Macrostomum sp.: These worms suck. Marine Biology, 145(2), 373– 380.

Simmons, L. W., & Siva-Jothy, M. T. (1998) Sperm competition in insects: Mechanisms and the potential for selection. Sperm Competition and Sexual Selection (pp. 341-434). Academic Press, London.

Simmons, L. W. (2001). Sperm competition and its evolutionary consequences in the insects. Princeton University Press.

Simmons, L. W., and Moore, A. J. (2009). Evolutionary quantitative genetics of sperm. In Sperm biology (pp. 405–434). Academic Press, London.

Snook, R.R. (2005). Sperm in competition: not playing by the numbers. Trends in ecology & evolution, 20(1), 46–53.

Stockley, P., Gage, M. J. G., Parker, G. A., & Møller, A. P. (1997). Sperm competition in fishes: the evolution of testis size and ejaculate characteristics. The American Naturalist, 149(5), 933–954.

Stoffel, M. A., Nakagawa, S., and Schielzeth, H. (2017). rptR: repeatability estimation and variance decomposition by generalized linear mixed-effects models. Methods in Ecology and Evolution, 8(11), 1639–1644.

Tourmente, M., Gomendio, M., and Roldan, E. R. (2011). Sperm competition and the evolution of sperm design in mammals. BMC evolutionary biology, 11(1), 1–10.

Vellnow, N., Vizoso, D. B., Viktorin, G., and Schärer, L. (2017). No evidence for strong cytonuclear conflict over sex allocation in a simultaneously hermaphroditic flatworm. BMC Evolutionary Biology, 17, 1-14.

Vellnow, N., Marie-Orleach, L., Zadesenets, K. S., & Schärer, L. (2018). Bigger testes increase paternity in a simultaneous hermaphrodite, independently of the sperm competition level. Journal of Evolutionary Biology, 31(2), 180–196.

Vizoso, D. B., Rieger, G., and Schärer, L. (2010). Goings-on inside a worm: Functional hypotheses derived from sexual conflict thinking. Biological Journal of the Linnean Society, 99(2), 370–383.

Willems, M., Leroux, F., Claeys, M., Boone, M., Mouton, S., Artois, T., and Borgonie, G. (2009). Ontogeny of the complex sperm in the macrostomid flatworm *Macrostomum lignano* (Macrostomorpha, Rhabditophora). Journal of Morphology, 270(2), 162–174.

Zadesenets, K. S., Vizoso, D. B., Schlatter, A., Konopatskaia, I. D., Berezikov, E., Schärer, L., and Rubtsov, N. B. (2016). Evidence for karyotype polymorphism in the free-living flatworm, *macrostomum lignano*, a model organism for evolutionary and developmental biology. PLoS ONE, 11(10), 1–24.

Zadesenets, K. S., Jetybayev, I. Y., Schärer, L., and Rubtsov, N. B. (2020). Genome and karyotype reorganization after whole genome duplication in free-living flatworms of the genus macrostomum. International Journal of Molecular Sciences, 21(2).

