## Supplementary material for "Sperm competition favours intermediate sperm size in a hermaphrodite": Supplemntary information

### Supplementary Information

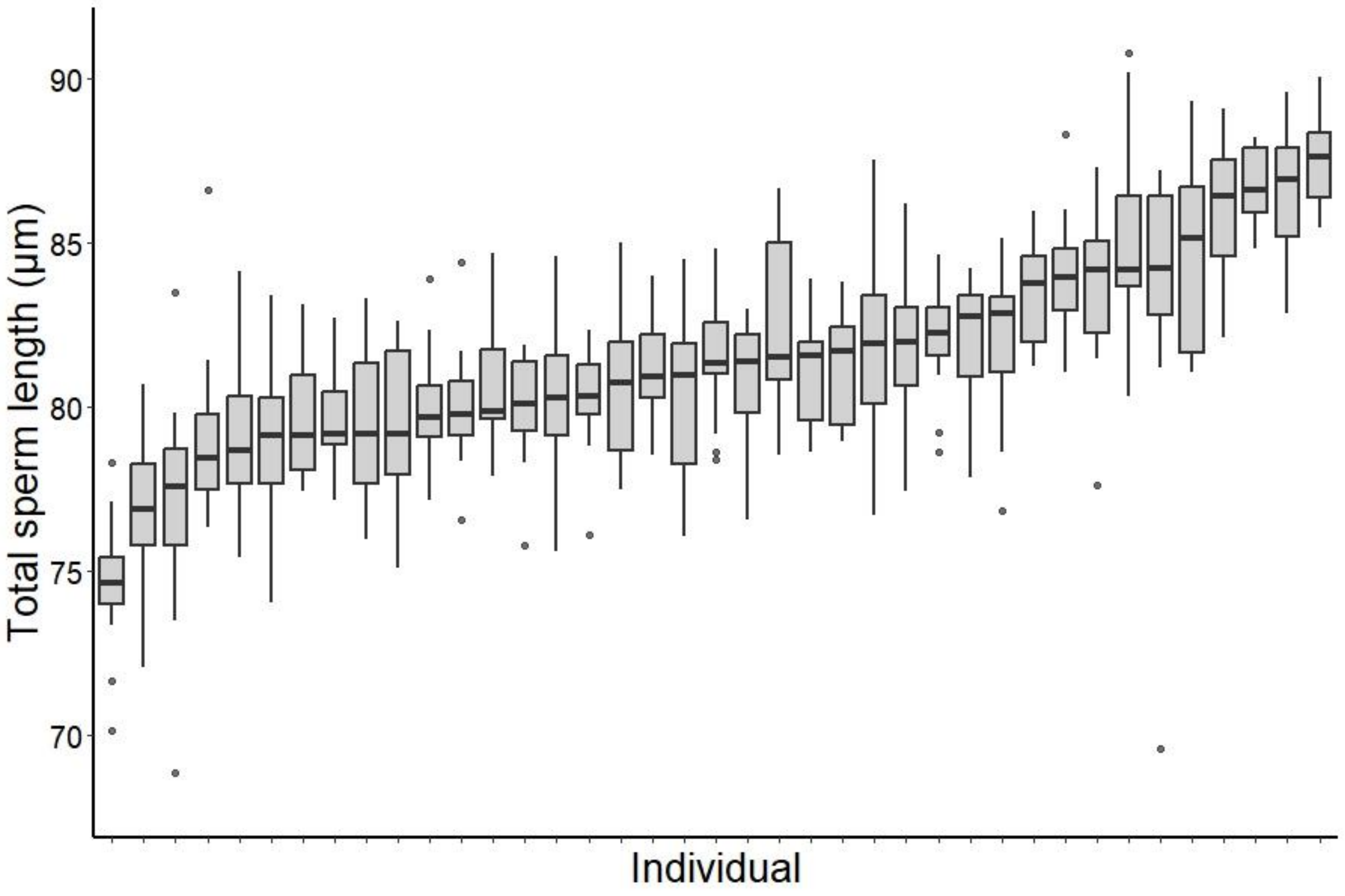


Fig S1. Between-individual variation in total sperm length in the outbred BAS1 population (n = 39). Individuals are arranged in increasing order of their median total sperm length (see Table S1 for statistical tests).

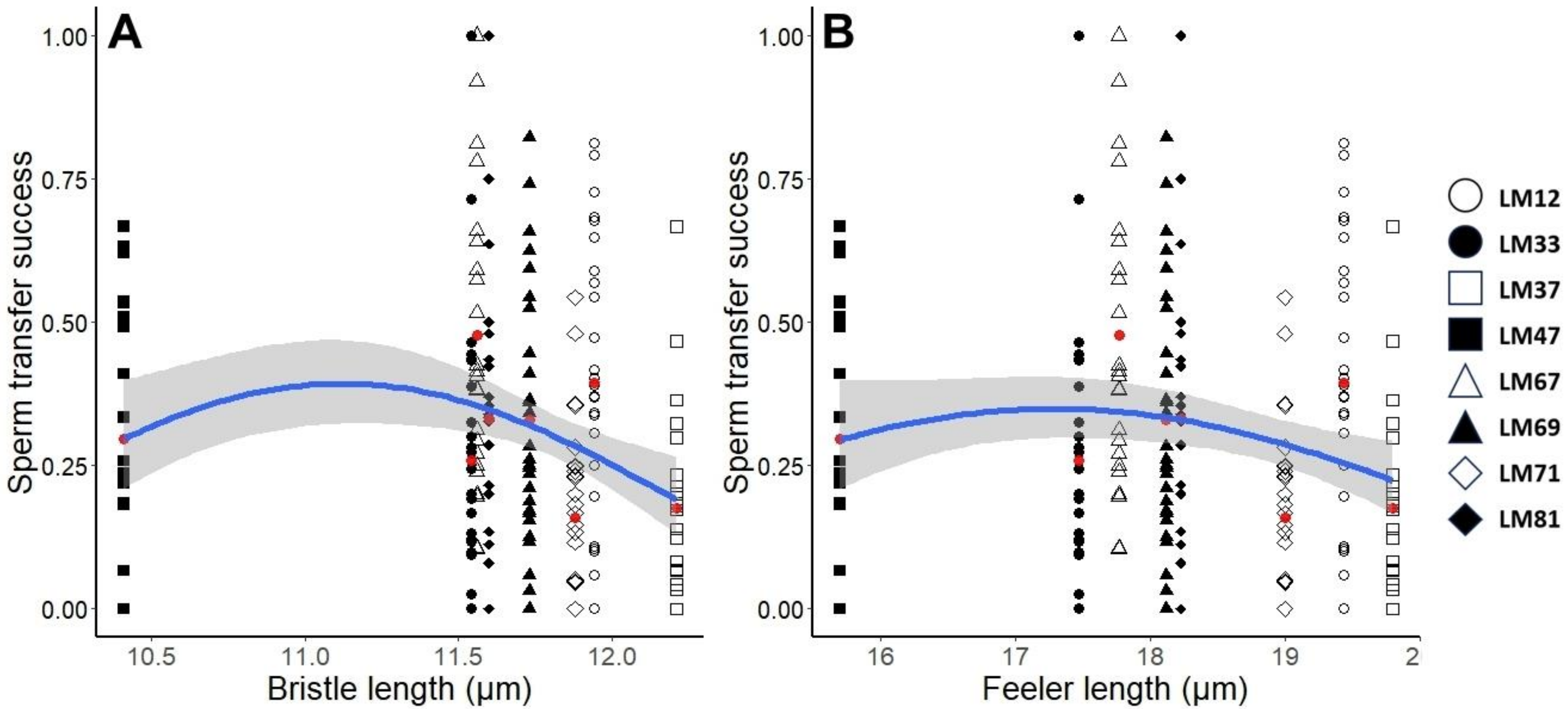


Fig S2. Relationship between sperm transfer success and (A) lateral bristle length and (B) anterior feeler length. Symbols indicate different LM lines. Red dots indicate mean sperm transfer success of each line. Solid blue line shows the fit of the quadratic model and grey regions indicate the 95% confidence intervals.

Table S1. Statistical tests and variation in sperm traits in the BAS1 culture. (based on all sperm cells measured, n = 591) Results of Kruskal-Wallis analysis of variance testing for variation in sperm morphology between individuals (n = 39). (SD -Standard deviation; SE - Standard error; CV - Total coefficient of variation; WCV - Mean within-ejaculate coefficient of variation)

| Sperm trait | Mean ± SD (µm) | Repeatability ± SE | CV (%) | WCV (%) | $X^2$ | *P* |
|---|---|---|---|---|---|---|
| Total sperm length | 79.4 ± 3.51 | 0.962 ± 0.03 | 4.31 | 2.65 | 347.5 | $<2.2e^{-16}$ |
| Feeler length | 16.92 ± 1.03 | 0.671 ± 0.167 | 6.10 | 4.87 | 202.54 | $<2.2e^{-16}$ |
| Bristle length | 10.91 ± 0.6 | 0.911 ± 0.065 | 5.52 | 4.30 | 231.42 | $<2.2e^{-16}$ |

Table S2. Overall phenotypic variance in sperm traits and the variance due to LM line identity and individual worm identity, from linear mixed models (Proportions in brackets).

| Sperm trait | Phenotypic Variance | Mixed model effects - variance | | |
|---|---|---|---|---|
| | | LM identity | Worm identity | Residual variance |
| Total sperm length | 38.259 | 18.292 (0.45) | 14.748 (0.37) | 7.201 (0.18) |
| Feeler length | 3.085 | 1.138 (0.36) | 0.999 (0.31) | 1.073 (0.33) |
| Bristle length | 0.723 | 0.178 (0.24) | 0.295 (0.40) | 0.270 (0.36) |

Table S3. Information on the cultures and inbred lines of *Macrostomum lignano*.

| Culture/ inbred line | Origin/ derived from | Method used | GFP+/GFP- | Use in this study | Reference |
|---|---|---|---|---|---|
| LS1 (outbred) | Field caught worms | Field sampling | GFP- | Not used | Marie-Orleach et al., 2013 |
| DV1 (inbred) | LS1 | Full- and half-sib crossing from pairs | GFP- | Common competitors in mating trials | Janicke et al., 2013; Vellnow et al., 2017 |
| HUB1 (inbred) | DV1 | Transgene (DNA) injection into single cell embryo | GFP+ | Not used | Janicke et al., 2013 |
| BAS1 (outbred) | LS1 | Backcrossing HUB1 | GFP+ | Measuring natural variation in sperm size | Marie-Orleach et al., 2016 |
| LM lines (inbred) | DV lines (similar to DV1) | Backcrossing HUB1 | GFP+ | Focal worms in mating trials | Marie-Orleach et al., 2017 |
